# Frontal and parietal alpha oscillations reflect attentional modulation of cross-modal matching

**DOI:** 10.1101/477034

**Authors:** Jonas Misselhorn, Uwe Friese, Andreas K. Engel

## Abstract

Multisensory perception is shaped by both attentional selection of relevant sensory inputs and exploitation of stimulus-driven factors that promote cross-modal binding. Underlying mechanisms of both top-down and bottom-up modulations have been linked to changes in alpha/gamma dynamics in primary sensory cortices and temporoparietal cortex. Accordingly, it has been proposed that alpha oscillations provide pulsed inhibition for gamma activity and thereby dynamically route cortical information flow. In this study, we employed a recently introduced multisensory paradigm incorporating both bottom-up and top-down aspects of cross-modal attention in an EEG study. The same trimodal stimuli were presented in two distinct attentional conditions, focused on visual-tactile or audio-visual components, for which cross-modal congruence of amplitude changes had to be evaluated. Neither top-down nor bottom-up cross-modal attention modulated alpha or gamma power in primary sensory cortices. Instead, we found alpha band effects in bilateral frontal and right parietal cortex. We propose that frontal alpha oscillations reflect the origin of top-down control regulating perceptual gains and that modulations of parietal alpha oscillations relates to intersensory re-orienting. Taken together, we suggest that the idea of selective cortical routing via alpha oscillations can be extended from sensory cortices to the frontoparietal attention network.

## Introduction

Human perception is governed by constant influx of information through multiple sensory channels. The act of perceiving routes information flow by active engagement with the multisensory environment, causing sensory inputs to be constantly shaped by modulatory signals reflecting behavioural goals, contextual demands and structural properties of the environment. Spotting a singing bird in a tree, for instance, does not depend on tactile processing but on evaluating visual and auditory signals for temporo-spatial congruence. Lacing shoes, on the other hand, makes little use of audition but integrates vision and tactile perception in a goal-directed manner. These examples illustrate that multisensory perception is shaped by top-down and bottom-up modulation of sensory inputs^1^. Attempts to understand multisensory perception accordingly need to address neural mechanisms underlying both selection of relevant sensory input and exploitation of stimulus-driven, intersensory similarities that promote cross-modal binding.

A well-described mechanism of attentional stimulus selection is gain regulation of population responses in sensory regions^2,3^. MEG and EEG studies have shown that gain regulation can be reflected in alpha band dynamics^4^. This has been demonstrated in several studies with respect to anticipatory biasing of visual processing in visuospatial cueing^5–8^. In these experiments, amplitude of alpha oscillations in the cued hemisphere is typically lower compared with the contralateral hemisphere, while gamma oscillations show the reverse pattern^9^. It was proposed that alpha oscillations might inhibit cortical processing in a cyclic manner delivering pulsed inhibition to gamma oscillations involved in local cortical processing^10^. Thus, information flow in cortex would at least partly be gated by inhibition of irrelevant nodes^11^. Although originally proposed for the visual system, the idea of gating by inhibition is not restricted to visual cortex but could in principle be applied to any cortical system. While there is some evidence that such alpha modulations are present for auditory^12^ and somatosensory^13^ modalities in spatial cueing tasks, a potential generalization to a larger number of cortical areas and tasks remains unclear.

With respect to stimulus-driven aspects of multisensory perception, a multitude of studies show behaviourally beneficial effects of concurrent and congruent multisensory stimuli^14–16^. While this is well established, underlying mechanisms remain largely elusive^17–19^. While some evidence suggests that input to distinct modalities is processed in parallel and only converges later in regions of the temporal and parietal lobe^20–22^, other evidence points to interactions already at the level of primary sensory regions^23–25^. The disparity of findings is not surprising given that influences driving cross-modal integration range from psychophysical (spatial/temporal congruence) to memory-dependent (semantic congruence and cross-modal correspondences) factors. Yet, a linking observation is that both low- and high-level integration have been associated with changes in gamma band activity^19,21,24,26,27^.

In a recent study on audio-visual matching, we demonstrated that top-down attention modulated alpha/gamma power in sensory regions while cross-modal congruence affected right temporal beta band coupling^28^. These findings are well in line with the literature reviewed above. In the present study, we employed an extension of our earlier experimental design now incorporating concurrent trimodal stimulation^16^. That is, we investigated whether the mechanisms outlined above would hold in a situation with strong sensory drive and cross-modal competition. We presented the same trimodal stimuli in two distinct attentional conditions (top-down influence), focused on either visual-tactile (VT) or audio-visual (AV) components, for which cross-modal congruence (bottom-up influence) of amplitude changes had to be evaluated (Fig. 1a). We expected top-down cross-modal attention to selectively enhance primary sensory alpha activity for irrelevant modalities and decrease alpha activity for attended modalities. This increase/decrease in alpha power might be accompanied by a decrease/increase in gamma band activity. As a bottom-up effect of cross-modal binding, beta/gamma band activity in sensory cortices or temporal/parietal cortex is expected to be modulated by cross-modal congruence.

**Figure 1:**
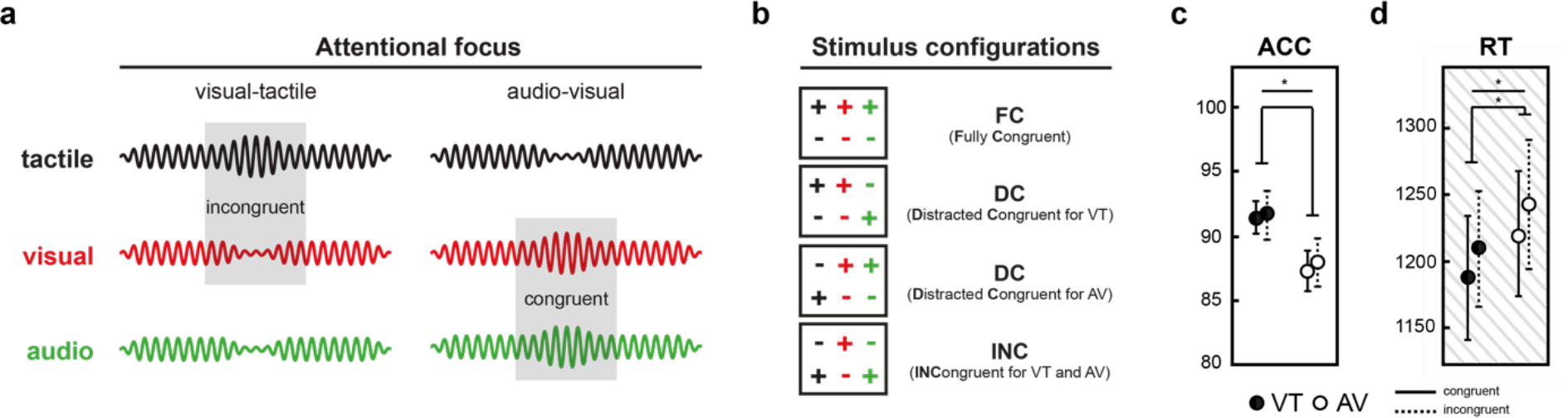
Schematic of matching task and behavioural results. **(a)** Illustration of example trials in the two attentional conditions. Under visual-tactile focus (left), the changes in the target stimuli are incongruent, as the tactile stimulus (black) increases in intensity while the visual stimulus (red) decreases. Under audio-visual focus (right), target change is congruent, since visual and auditory (green) stimuli both increase in intensity. **(b)** In each block, all possible stimulus configurations occurred with equal probability. Intensity changes are depicted by coloured plus/minus signs (colour coding as introduced in a). **(c)** Accuracy (ACC) in percentage correct. Error bars represent standard deviations. *ATTENTION* significantly affected accuracy of responses. **(d)** Reactions times in milliseconds (RTs). Error bars represent standard deviations. *ATTENTION* as well as *CONGRUENCE* significantly affected the timing of responses (Note: RTs were collected in our previous behavioural study16 from the same sample of participants).

## Results

### Psychophysics, questionnaire and behaviour

The trimodal stimulus material was designed such that the target amplitude changes in each modality were equally salient. This was achieved by estimating detection thresholds for each modality and change direction separately using a psychophysical staircase procedure^29^. Yet, a questionnaire that was completed during debriefing of the preceding behavioural study^16^ indicated that subjective salience of the sensory components when presented jointly was in fact not equal, but strongest for the visual component. In particular, participants reported that the visual component was hardest to ignore when it was task irrelevant (Wilcoxon rank sum test, Bonferroni correction; average ranks: V=1.4, A=2.3, T=2.1; V-T: *p*_corr_ = 0.0074, V-A: *p*_corr_ = 0.7902, T-A: *p*_corr_ = 0.7200). Conversely, visual change direction was reported to be easiest to classify compared with both other modalities (average ranks: V=1.6, A=2.0, T=2.4; V-T: *p*_corr_ = 0.0135, V-A: *p*_corr_ = 0.0032, T-A: *p*_corr_ = 3.2427). Anecdotally, many participants reported that their strategy was to inhibit the respective dominant modality to increase attentional focus on the relatively weaker stimulus. Overall, participants rated VT condition as easier than AV condition (average ranks: VT=1.3, AV=2.0; VT-AV: *p*_corr_ = 0.0171) and AV in turn as easier than AT (average rank: AT=2.7; VT-AT: *p*_corr_ = 0.0061, AV-AT: *p*_corr_ = 0.0171).

The accuracy of responding was analysed with a repeated measure analysis of variance (ANOVA) with factors *ATTENTION* (VT/AV) and *CONGRUENCE* (congruent/incongruent). When participants attended VT, accuracy of responding was significantly higher compared with the AV condition (ACC; *p* = .005, 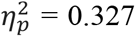). No interaction effects between *ATTENTION* and *CONGRUENCE* were observed. In order to minimize muscle artifacts in the EEG, participants gave verbal responses only after stimulus offset. We thus present data on response timing by re-analysis of data collected in the preceding behavioural study^16^ from the same sample of participants with the statistical model explained above. Responses to cross-modally congruent pairs were faster (Fig. 1d; *p* < .001, 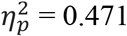), as well as responses to VT attention trials (*p* = .036, 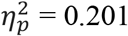). There was no interaction between the main effects.

### ROI analysis

In Figure 2, we present an overview of time-frequency dynamics during the task as well as distributions of band-limited power for theta, alpha, beta and gamma bands in source space. Frequency bands of interest were defined individually based on visual inspection of each participant’s time-frequency plot (see *Methods* for details). Statistical analysis was focused on the interval containing the changes in stimulus intensity ([0; 300] ms relative to change onset; cf. Fig. 2a top panel). First, we investigated changes in oscillatory power relative to a baseline ([−400; −100] ms relative to stimulus onset) for regions of interest (ROI) in visual, auditory and somatosensory cortices (Fig. 3a). To that end, baseline-corrected power was computed in source-level EEG data, averaged for the whole change epoch and subjected to a repeated measures ANOVA with factors *ROI*, *ATTENTION* and *CONGRUENCE*, separately for each frequency band (Fig. 3). Significant effects of *ROI* were observed for theta, alpha and beta bands corresponding to the cortical topographies depicted in Figure 2 (for all, *p* < .001 and 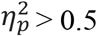; see Fig. 3b). In the gamma band, *ROI* did not explain a significant amount of variance (*p* = .689). Simple effects analysis for the lower frequency bands showed that decreases in power were significantly stronger in visual compared with both auditory and somatosensory ROIs (for all comparisons, *p* < .001; Fig. 3b). Power changes in auditory and somatosensory ROIs did not differ (for all, *p* > .05). The main effects and interactions of *ATTENTION* and *CONGRUENCE* were not significant (for all, *p* > .05; Fig. 3c+d).

**Figure 2:**
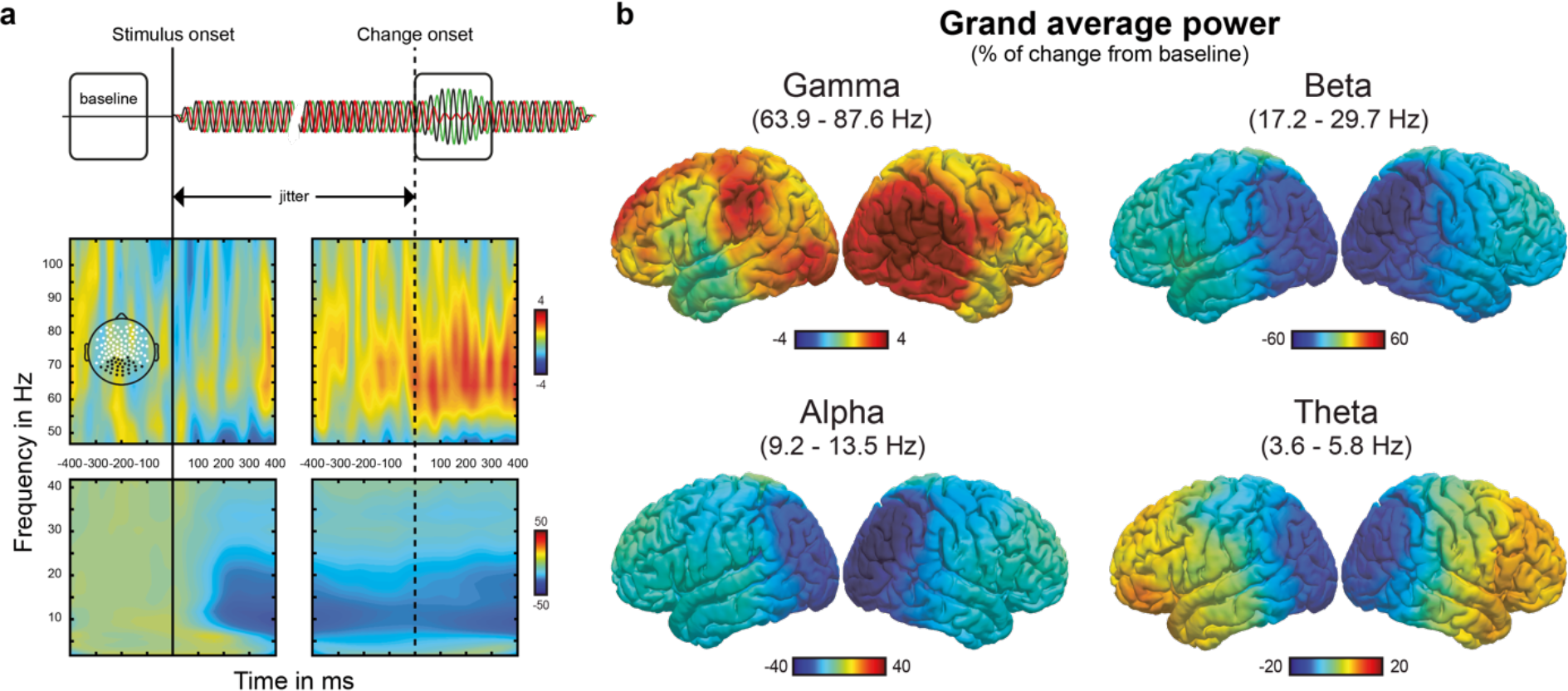
Average time-frequency dynamics in sensor and source space. **(a)** Time-frequency dynamics in sensor space at occipito-parietal channels (topography shown in upper left panel), time locked to stimulus onset (left panels, black solid vertical line) and change onset (right panels, black dashed vertical line). ***Top:*** Schematic of temporal trial structure displaying the two relevant time-windows used for analysis: a baseline ([−400; −100] ms relative to stimulus onset) separated by a jitter (stimulus to change onset between 700 and 1000 ms) from the change interval ([0; 300] ms relative to change onset). ***Center, bottom:*** Time-frequency plots from posterior sensors. Values represent percentage of change from baseline. **(b)** Distribution of band-limited power on the cortical surface in the change interval relative to baseline.

**Figure 3:**
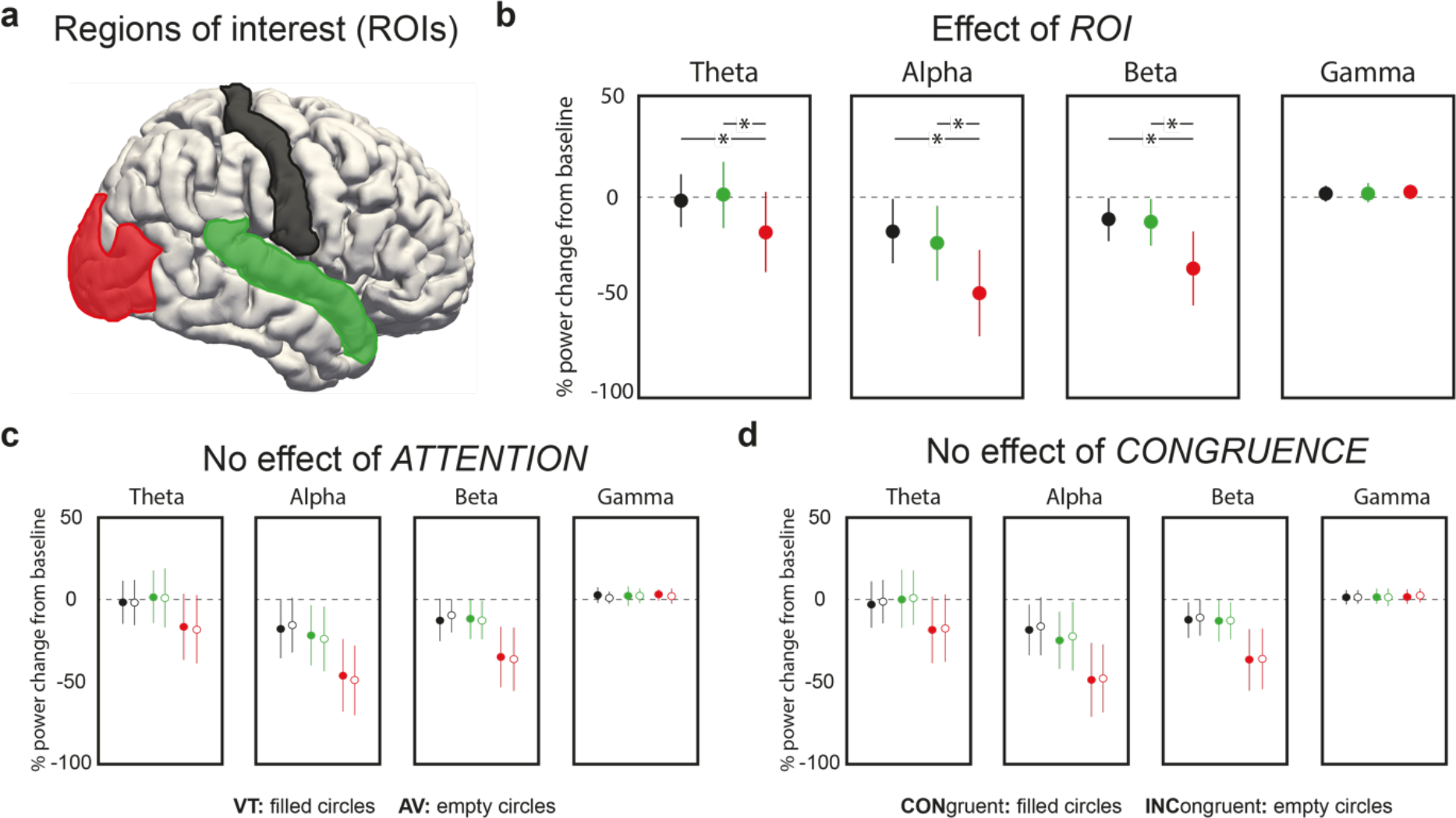
Regions of interest (ROI) analysis of band-limited power during change epoch in primary sensory areas. **(a)** Primary sensory areas used for ROI ANOVA with factors *ATTENTION* (VT vs. AV), *CONGRUENCE* (congruent vs. incongruent) and *ROI* (visual vs. auditory vs. somatosensory); visual = red, auditory = green, somatosensory = black. **(b)** Effect of *ROI* is significant for theta, alpha and beta bands, but not gamma band. Asterisk signifies significant comparisons (*p* < .001). **(c)** No significant effect of *ATTENTION*. **(d)** No significant effect of *CONGRUENCE*.

### Cluster statistics

To complement the ROI analysis, we conducted a whole-brain analysis of task-related power changes in the change interval ([0; 300] ms relative to change onset). Multiple comparisons in space were controlled by means of nonparametric cluster-based permutation statistics (see *Methods* for details). Multiple testing due to the factorial design and the four frequency bands was Bonferroni-corrected (12 comparisons, α_crit_ = 0.05/12 = .0042). We report uncorrected *p*-values of all clusters significant at α_crit_.

For the *ATTENTION* contrast (VT *minus* AV), clusters of significant differences were found in the alpha band. In two roughly symmetric frontocentral clusters, VT attention was associated with relatively lower alpha power when compared with AV attention (Fig. 4a). In the left hemisphere, the cluster was situated in the border region of pre-central gyrus, middle and superior frontal gyrus encompassing the frontal eye fields (FEF; *p* = 0.0026). In the right hemisphere, the cluster was situated similarly but expanded further into pre- and post-central gyrus (*p* = 0.00043). In order to investigate whether this attentional alpha modulation occurred only during the presentation of relevant stimulus features, or whether it would possibly resemble the spatial cueing effects that are most prominent prior to stimulus presentation, we analysed time courses of alpha power modulations within these two clusters (Fig. 4b, see *Methods*). Descriptively, alpha power was lower for VT than for AV attention throughout the entire time course. This difference was significant before stimulus onset in both hemispheres (left: [−400; −312] ms, right: [−165; 43] ms), in a short epoch prior to change onset in the left hemisphere ([−348; −286] ms) and throughout the whole change epoch in both hemispheres (left: [−69; 177] ms and [211; 400] ms, right: [43; 400] ms). In order to further elucidate this effect, we split the group according to responses to the questionnaire and computed the same time courses as before. While only small, but consistent *ATTENTION* differences were seen for the group that reported having difficulties with ignoring visual input, larger differences were seen for the group having less difficulty ignoring visual input (Fig. 4c).

**Figure 4:**
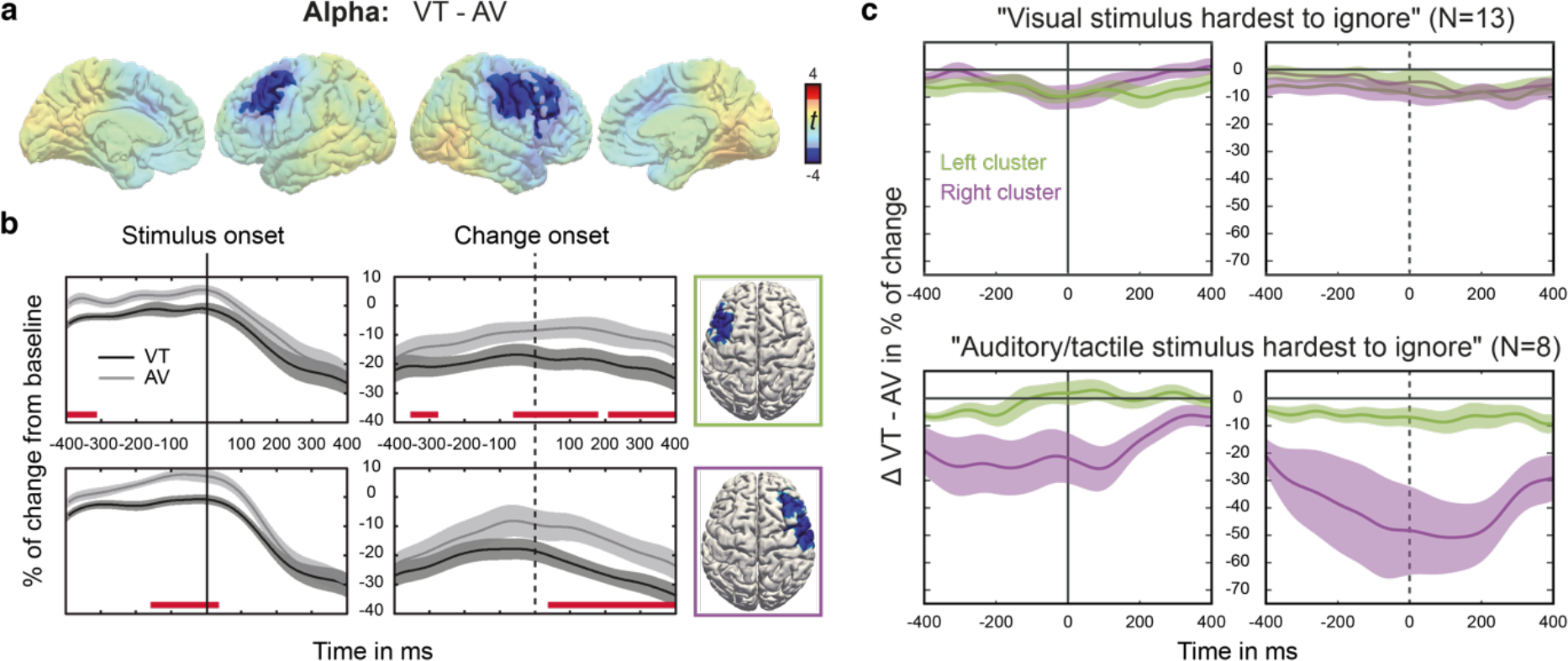
Effect of *ATTENTION*. **(a)** Cluster-based permutation statistic of the contrast VT *minus* AV in the alpha band during change interval. T-values of the respective contrasts are depicted. Shaded voxels are non-significant at cluster-level. **(b)** Time-course of alpha power within significant clusters. Shading corresponds to the standard error of the mean. Red bars indicate temporal regions of significance as determined by permutation testing (see *Methods* for details). **(c)** *ATTENTION* difference (VT-AV) of alpha power in left (green) and right (purple) cluster depicted for sub-groups. These sub-groups were formed according to responses to a questionnaire completed after participation in our preceding behavioral study16.

When evaluating the effect of *CONGRUENCE* (attended congruent *minus* attended incongruent), clusters of significant differences were found in the alpha and theta bands (Fig. 5). Descriptively, theta power was higher for congruent trials compared to incongruent trials in large parts of the medial aspect of both hemispheres (Fig. 5a). This effect was significant in a left hemispheric cluster extending from posterior to anterior cingulate cortex (*p* = 0.00247). In a next step, we analysed the differential contributions of fully congruent (FC) and distracted congruent (DC) trials to the overall effect of congruence. Those two sub-conditions were expected to differ with respect to their neurophysiological underpinnings because FC trials contained no conflicting inputs, while DC trials did. Thus, we computed and qualitatively compared the contrasts between FC respectively DC trials and incongruent trials. Cortical maps represent uncorrected *p*-values shaded for values above 0.05 (Fig. 5c,e; see *Methods* for details). By comparing these contrasts, it is noticeable that FC contributed most strongly to the overall theta difference in cingulate cortex (Fig. 5c). There, theta power was higher in cingulate cortex of both hemispheres, whereas no marked differences can be seen when comparing DC and incongruent trials (Fig. 5e).

**Figure 5.**
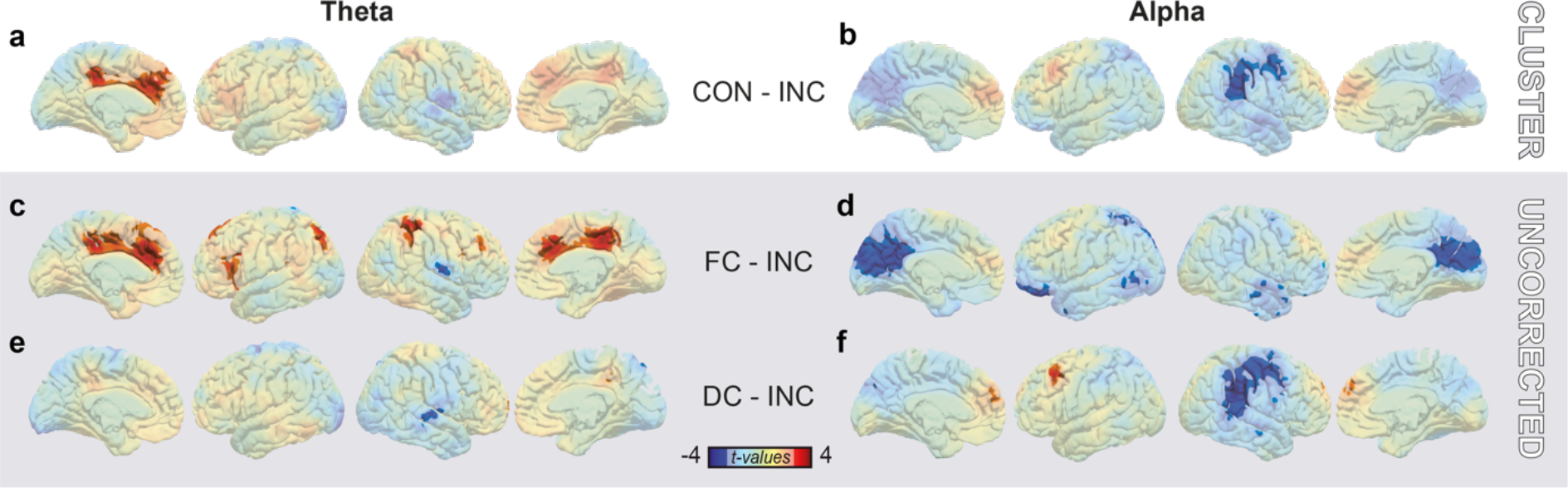
Effect of *CONGRUENCE*. **(a+b)** Cluster-based permutation statistic of the overall *CONGRUENCE* contrast in the theta (a) and alpha (b) band during change interval. All subplots depict *t*-values. Shaded voxels are non-significant as determined by cluster statistic. **(c-f)** Post-hoc comparisons of sub-condition contrasts. Presented statistical maps are uncorrected, voxels with *p*-values above 0.05 are shaded. **(c)** Contrast between theta power in fully congruent (FC) and incongruent trials. **(d)** Contrast between alpha power in fully congruent (FC) and incongruent trials. **(e)** Contrast between theta power in distracted congruent (DC) and incongruent trials. **(f)** Contrast between alpha power in distracted congruent (DC) and incongruent trials.

In the alpha band, cross-modal congruence modulated power in large parts of medial occipital and lateral parietal/frontal cortex. Descriptively, bilateral medial superior frontal cortex and left MFG showed positive *CONGRUENCE* differences (Fig. 5b), that is, stronger decrements in alpha power for incongruent trials (note that overall alpha power decreased in these regions, cf. Fig. 2b). These, however, were not significant at the cluster-level. Negative *CONGRUENCE* differences, that is, stronger decrements in alpha power for congruent trials can be seen in bilateral medial occipito-parietal cortex and right parietal/central cortex (Fig. 5b). While differences in medial-occipital cortex barely missed significance at the cluster-level (*p* = 0.0044 > 0.0042 = α_crit_), a cluster covering the right temporoparietal junction (TPJ), but stretching rostrally towards right MFG, was significant (*p* = 0.00247). Next, we disentangled contributions from FC and DC trials as before for theta power. While FC trials seemed to drive the *CONGRUENCE* difference in medial occipito-parietal cortex (Fig. 5d), DC trials contributed strongly to the effect in right TPJ (Fig. 5f).

## Discussion

We investigated bottom-up and top-down modulation of sensory processing in a cross-modal matching task involving visual, auditory and tactile perception. Contrary to our expectations, we did not find alpha/gamma oscillations in primary sensory areas to be modulated by bottom-up or top-down cross-modal attention. This finding is surprising given that both processes have often been noted to be accompanied by alpha/gamma modulations in sensory cortices^5,9,12,13,24,28,30^. Our explanation for the lack of primary sensory modulation is the nature of the task: many, if not most, multisensory studies employ detection tasks with near-threshold sensory stimulation. In these situations of low sensory drive, both bottom-up and top-down modulation of sensory input can be expected to have higher impact compared with situations of strong sensory drive. Stimulus-driven cross-modal enhancement by spatio-temporal congruence, for instance, is assumed to obey the law of inverse effectiveness, meaning that there is an inverse relationship between possible cross-modal enhancement and stimulus intensity^31^. Here, however, all stimulus intensities were clearly supra-threshold with superimposed amplitude changes. Top-down as well as bottom-up modulation of responses in sensory cortices might thus be subtle and hence not detectable by EEG. Instead, we found theta oscillations in cingulate cortex and most notably alpha oscillations in frontal and parietal cortex to be modulated. In the following, we propose that frontal alpha oscillations might reflect the origin of top-down control regulating perceptual gains and that parietal alpha oscillations could relate to intersensory (re-)orienting. Theta activity in cingulate cortex is finally discussed in the context of adaptive task-switching.

Task-related reduction of alpha power in bilateral FEF and MFG as well as in right pre- and post-central gyrus was stronger for VT attention compared to AV attention. In a time-resolved post-hoc analysis, we showed that this difference was significant even before stimulus onset. Besides their role in oculomotor control, the FEF have been described as important structures in top-down attention^32^. In a study using TMS, Grosbras and Paus^33^ showed that disruption of activity in right FEF shortly before the onset of the target in a visuospatial covert attention task facilitated responses. Conversely, 10 Hz TMS over the right FEF was shown to impair visual search of unpredictable items with low salience^34^. Moore and Armstrong reconciled this conflicting evidence by suggesting that the FEF may have a general role in regulating visual gain^35^. In their study, electric stimulation of FEF in the monkey either enhanced or inhibited responses to visual stimuli in V4 depending on whether retinopically corresponding sites were stimulated. Studies in humans supported this idea by showing that TMS over FEF could increase phosphene or contrast sensitivity of extrastriate cortex^36,37^. This top-down modulation of visual cortex was demonstrated even in the absence of sensory input^38^. Likewise, anticipatory alpha and stimulus-related gamma activity in occipito-parietal cortex could be modulated by TMS over FEF^8^. The strength of modulation was shown to correlate with the strength of structural connectivity between frontal and parietal cortex via the superior longitudinal fasciculus^39^. Thus, animal and human studies jointly conclude that the FEF can dynamically modulate the gain of up-stream visual cortex independent of sensory input. In our study, FEF/MFG is proposed to have facilitated cross-modal matching by modulating visual gain to counter visual dominance. Although stimulus intensity was titrated to be balanced across modalities (see *Methods*), we have reason to assume that perceived salience was highest for the visual component. In a questionnaire that was completed during debriefing of the preceding behavioural study, we asked participants to rank the difficulty to ignore a given modality. Most participants reported that the visual component was hardest to ignore. Conversely, most participants ranked the visual intensity change as the easiest to classify as either increase or decrease (V > A > T). This finding is in line with a pattern of sensory dominance found for combinations of visual, auditory and somatosensory stimuli in a discrimination task^40^. Sensory dominance can be problematic under the assumption that cross-modal matching is not independent of perceptual gain. This is most likely the case for stimulus-driven aspects of multisensory integration – the idea of inverse effectiveness, after all, assumes multimodal stimuli of low but comparable intensity^31^. Anecdotally, our participants reported that they strategically supressed the subjectively stronger stimulus in order to improve focus on the subjectively weaker stimulus. Consequentially, the need for adjustment of sensory dominance should be higher for attended VT compared with attended AV condition because subjective salience differed more between V and T than between V and A. Next, we used the questionnaire data to form sub-groups that either reported to have had particular difficulty in supressing visual stimuli or not. Although both groups showed time courses of VT-AV differences comparable to that seen in the whole group, stronger differences were seen for the group with less difficulty suppressing visual input, most pronounced in the right hemispheric cluster. This finding could point to a lateralized role of the right FEF in attentional modulation of bilateral sensory cortex, as has been suggested before^33^. Taken together, we speculate that decreased power of alpha oscillations in FEF/MFG might relate to down-regulation of visual gain, which might have been more pronounced in VT conditions to counter-act unequal subjective salience of the stimuli to be matched.

As discussed above, balancing perceptual gains across modalities by top-down modulation likely enables optimal use of stimulus-driven aspects of cross-modal matching. These bottom-up factors were ubiquitous in this task; on each trial, participants were simultaneously confronted with three salient events, that is, intensity changes in each modality. Although each change of intensity by itself possessed some degree of bottom-up salience, we suggest that cross-modal congruence amplified salience through cross-modal binding^19^. When cross-modal binding was enhanced between attended modalities, responses were facilitated. This was especially pronounced for fully congruent trials where conflict, and thus the need for actual matching, was absent^16^. All other trials were either distracted congruent (attended modalities changed congruently, but the distractor diverged) or attended incongruent (one of the attended modalities was congruent to the distractor). In these cases, cross-modal binding was always stronger between two given modalities compared to the respective third. When contrasting the EEG of these trials, we find alpha band effects in the right temporoparietal junction (rTPJ) and right MFG. Specifically, distracted congruent conditions were associated with enhanced decreases of alpha power in these regions compared with attended incongruent trials. The TPJ receives inputs from visual, auditory and somatosensory cortex and is richly connected to temporal and frontal sites, making it an important hub for the interaction of multisensory integration and attention^41^. Accordingly, lesions to the right TPJ typically result in neglect^42,43^. A dominant interpretation of rTPJ‘s functional role is its involvement in spatial re-orienting based on stimulus salience^44^. In a model integrating goal-directed and stimulus-driven attention, it has been suggested that a dorsal network comprising FEF and IPS instantiates attentional sets. As a counterpart, a ventral network comprising rTPJ and right ventral frontal gyrus mediates bottom-up signals acting as a circuit-breaker for the dorsal system^44^. Studies employing multisensory paradigms have noted rTPJ’s involvement in processing cross-modal congruence. In a study investigating visual-tactile pattern matching, pre-stimulus alpha and beta power in right supramarginal gyrus differentiated between detection and congruence-evaluation tasks^45^. Another study showed that alpha power in right posterior regions was more strongly supressed during congruent compared with incongruent audio-visual speech presentations^46^. Taken together, we suggest that the rTPJ detects the increased salience of congruent cross-modal events. While in each trial attention might, in principle, be captured by any of the three modalities, cross-modal binding by congruence might serve as a reliable “cue” for re-orienting towards the relevant modalities. Thereby, cross-modal binding between attended modalities might support intersensory re-orienting. We suggest that enhanced power decreases in rTPJ during congruent trials reflects binding facilitating this intersensory attentional capture.

As pointed out above, fully congruent trials were characterised by the absence of cross-modal conflict. In the EEG, these highly salient trials were associated with stronger alpha power reductions in medial occipito-parietal cortex when compared to incongruent trials (note, however, that this difference barely missed significance after Bonferroni correction). In an event-related potentials study featuring visual, auditory and somatosensory stimuli, RT facilitation was correlated with the latency of the P300, which was localised in precuneus^47^. Other research suggests that alpha power reductions in occipito-parietal cortex and P300 dynamics are functionally coupled^48^. Here, enhanced involvement of medial occipito-parietal cortex is proposed to reflect increased bottom-up salience due to multisensory enhancement, i.e., increased perceptual gains of concurrent congruent sensory input to more than one modality. In addition to their bottom-up sensory salience, fully congruent stimuli occurred in only 25 % of all trials and were thereby salient. Actual cross-modal matching was required only in the remaining 75 % of trials where two modalities changed congruently while the third modality diverged. An efficient strategy would accordingly be to “switch” between these two tasks, i.e., between detecting highly salient events and cross-modal matching of conflicting input. In addition to the alpha band effect in precuneus, fully congruent trials were also associated with a relative increase in theta power in bilateral cingulate cortex. Theta band activity in cingulate cortex has previously been related to the adjustment of stimulus response mappings^49^. Together with insular cortex, cingulate cortex is part of a salience network which has importance for both bottom-up detection of salient events and switching between large-scale networks to adaptively control behaviour^50^. Here, it is suggested that reduced alpha power in medial occipito-parietal cortex related to multisensory enhancement acts as a salience signal detected by cingulate cortex which in turn initiates adaptive task-switching behaviour.

The interpretation of the current results is limited in some important ways. First, we were not able to analyse response times which showed clear effects in our preceding behavioural study16. This was due to a delayed response interval where verbal responses were given, chosen in order to minimize muscular artifacts in the EEG. Second, accuracies were in general high and did not show congruence-related effects. Therefore, we could not test the behavioural relevance of our findings directly. Instead, we used questionnaire data to repeat analysis on sub-groups showing distinct behavioural strategies. The results of these post-hoc analyses are consistent with our interpretations.

Taken together, we provide evidence that cross-modal matching in complex multisensory environments relies on mechanisms of attention. Our results contrast with the majority of studies on multisensory integration concerned with stimulus detection where attentional load is typically low. Here, participants were confronted with a highly challenging multisensory setting. We speculate that, in order to counter the bias imposed by visual dominance, top-down regulation of gain in visual cortex supported an optimal exploitation of cross-modal similarities that promote perceptual binding. This was associated with decreased alpha band power in frontal cortices proposed to reflect the origin of top-down modulation. Likewise, bottom-up drive for cross-modal binding was related to changes in alpha power in right parietal cortex proposed to represent the bottom-up modulatory signal underlying intersensory re-orienting. Both findings provide evidence for an extension of the idea that alpha band dynamics indicate selective cortical routing beyond sensory cortex to the frontoparietal attention network.

## Methods

### Participants

Twenty-one participants entered the study and received monetary compensation for their participation. They were on average 23.8 ± 2.5 years old and 11 of them were female (10 male). Vision, audition and tactile perception were normal and none of them had a history of neurological or psychiatric disorders. After an explanation of the experimental procedure, participants gave informed consent. The ethics committee of the Hamburg Medical Association (Ärztekammer Hamburg) approved the study which was carried out in accordance with the declaration of Helsinki.

### Experimental procedure

We re-invited participants from a preceding behavioural pilot^16^ study to ensure reasonable performance on the task. While preparing EEG recordings, participants underwent a psychometric thresholding procedure for the stimulus materials used in the experimental blocks (see below; ~45 min). After the completion of EEG preparation and psychometric thresholding, participants completed at least three short training blocks to ensure stable performance on the task (~15 min). In total, subjects completed 10 experimental blocks (each ~15 min) with short breaks after every second block (~5 min). In total, experimental sessions took up to 4 hours. In order to collect sufficient data for statistical analysis, this procedure was carried out on two separate days with identical procedure for each participant.

### Experimental design

On each trial, we presented a trimodal stimulus set consisting of a visual, an auditory and a tactile component that each underwent a brief increase or decrease in intensity (see *Stimulus Material* for details). Block-wise, participants attended either visual-tactile (VT) or audio-visual (AV) bimodal pairs and ignored the respective third component. The task was to decide whether the attended bimodal pairs changed congruently (both attended stimuli increased or decreased in intensity) or incongruently (one attended stimulus increased and the other decreased in intensity; see Fig.1a). Participants responded verbally with “equal” (German: “gleich”) to congruent trials and “different” (German: “verschieden”) to incongruent trials. Verbal responses had to be withheld until stimulus offset to minimise myogenic artifacts. Therefore, reaction times (RTs) could not be evaluated. Instead, we present RT data of the same sample of participants from the behavioural study preceding the EEG study^16^. In each block, all eight possible stimulus configurations of increases and decreases across modalities were presented with equal probability (Fig. 1b). VT and AV blocks containing 64 trials presented in randomised order were alternating. Data were collected on two separate days with identical experimental procedure so that EEG data of 1280 trials was collected from each participant. Prior to statistical analysis, trials were pooled without taking change direction into account. For instance, fully congruent trials were both trials where all modalities underwent decrements and trials where all modalities underwent increments in intensity (Fig. 1b, pooling is indicated by boxes).

### Stimulus material

Visual contrast, auditory loudness and vibration strength were experimentally increased or decreased. The magnitudes of change per modality and direction were individually estimated prior to the experimental sessions using the same psychometric step function (QUEST) as described in Misselhorn et al.^16^. Intensity changes had a duration of 300 ms and onsets were jittered across trials between 700 and 1000 ms after stimulus onset (Fig. 2a). In total, sensory stimulation had a fixed duration of 2 s. As visual stimulation, an expanding circular grating was centrally presented against a grey background on a CRT screen with a visual angle of 5°. The auditory component consisted of a complex sinusoidal tone (13 sine waves: 64 Hz and its first 6 harmonics as well as 91 Hz and its first 5 harmonics, low-frequency modulator: 0.8 Hz) played back with audiometric insert earphones binaurally at 70 dB (E-A-RTONE 3A, 3M, USA). For tactile stimulation, high-frequency vibrations (250 Hz on C2 tactors, Engineering Acoustics Inc., USA) were delivered to the tips of both index fingers.

### EEG

EEG was recorded from 128 active electrodes (Easy Cap, Germany) including four ocular electrodes referenced to the nose. Data was sampled at 1000 Hz with an amplitude resolution of 0.1 µV using BRAINAMP MR amplifier (Brain Products, Germany) and digitised after analogue filtering (low cutoff: 10 s, high cutoff: 1000 Hz). Offline, data was down-sampled to 500 Hz and digitally filtered using the pop_eegfiltnew.m function provided by EEGLAB^51^ with default settings (high-pass: 1 Hz, low-pass: 120 Hz, notch: 49-51 Hz, 99-101 Hz). Epochs of 2.5 s were cut from −500 ms relative to stimulus onset until stimulus offset and normalised to the pre-stimulus baseline. Next, data was re-referenced to the common average and linear trends were removed from all epochs. From the four ocular channels, two bipolar channels for horizontal and vertical eye movements were derived.

#### Pre-processing

Trials with incorrect answer and large non-stereotypical artifacts were excluded from further processing. Subsequently, independent component analysis (ICA) was performed separately for low and high frequency bands (low band: 1-30 Hz, high band: 30-120 Hz). Thereby, stereotypical low-frequency artifacts (for instance eye movements and heart beat) and high-frequency artifacts (i.e. myogenic activity) could be separated more reliably from neuronal activity. For both bands, principal components analysis was performed first to reduce data such that 99 % of variance is retained. Subsequently, ICA was performed on the rank-reduced data using the infomax algorithm^51^. Artifactual ICs were identified and rejected with respect to time course, spectrum and sensor topography^52^. For the high band, saccade-related transient potentials were removed additionally^53^. Finally, all epochs were visually inspected and epochs with remaining artifacts were rejected. Furthermore, a subset of 19 electrodes (i.e. most outer facial, temporal and neck electrodes) was excluded from further analysis due to poor signal-to-noise ratio. Lastly, data was stratified such that all conditions in the ensuing analysis hold the same amount of data within subjects. On average, 426 ± 89 trials per participant entered the analysis.

#### Source reconstruction of band-limited signals

Cleaned data in low and high frequency bands were combined and epoched with respect to stimulus onset as well as change onset. Prior to filtering data into narrow bands by means of wavelet analysis, event related potentials were subtracted in order to remove phase-locked responses. A family of 40 complex Morlet wavelets 𝑤 with lengths of 2 s was constructed for logarithmically spaced frequencies between 2 and 120 Hz.

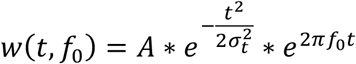

The number of cycles per wavelet (*m*) were logarithmically spaced between 3 and 10 and subsequently rounded off. Wavelets were normalised by factor 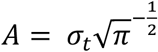 with *σ*_*t*_ = *m*/2*πf*_0_). Single trial data was convolved with the Morlet wavelets by multiplication in the frequency domain using fast fourier transformation with boxcar windows. Wavelet filtered single trial data was then reconstructed in source space using exact low-resolution brain electromagnetic tomography (eLORETA; regularisation: 0.05)^54^. Lead fields were computed for a three-shell head model^55^. The customised cortical grid was derived from a cortical surface provided by Freesurfer^56^ in MNI space by reducing the number of cortical nodes from 270000 to 10000. Dipole directions at each node of the cortical grid were estimated by means of singular value decomposition of the trial averaged spectral power individually for all bands and kept constant for all trials of the given participant. Power within each frequency bin was computed by multiplication with the complex conjugate (conj.m). Since we subtracted event-related potentials, we refer to the estimates as induced power. Average power in a time window before stimulus onset ([−400; −100] ms relative to stimulus onset) was used for baseline normalization. That is, the baseline was subtracted from all power estimates within each frequency bin and subsequently the difference was divided by the baseline and multiplied by 100. Thus, we computed percentage of change relative to baseline. By visual inspection of the resulting time-frequency landscapes, frequency bands in the theta, alpha, and beta range were chosen individually for each participant while gamma band was chosen uniformly based on canonical band width (mean values and range in parentheses; theta: 4.7 [3.6; 5.8] Hz, alpha: 11.5 [9.2; 13.5] Hz, beta: 23.0 [17.2; 29.7] Hz, gamma: 78.9 [63.9; 87.6] Hz). For that purpose, we computed individual time-frequency landscapes equivalent to the illustration of the grand average in Figure 2a for occipital, parietal, temporal and frontal regions separately. Spectral peaks/troughs of marked changes of spectral activity from baseline were detected and narrow frequency bands around these peaks/troughs were constructed. When clear peaks/troughs were missing, we resorted to canonical definitions of band-widths and oriented the boundaries of bands according to neighbouring bands. Thereby, we ensured that functionally relevant frequency bands were chosen.

### Statistical analysis

#### Behaviour

Accuracy of responding (ACC) within experimental conditions was analysed using a repeated-measures analysis of variance (ANOVA) with factors *ATTENTION* (VT/AV) and *CONGRUENCE* (congruent/incongruent). Greenhouse-Geisser correction was applied where necessary and effect sizes are reported as partial eta squared 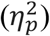. The timing of verbal responses was not analysed because subjects were instructed to withhold responses until stimulus offset. Instead, data from the previous behavioural study was re-analysed for the sub-sample of participants enrolled in this EEG study^16^. The same ANOVA as described above for ACCs was evaluated.

#### Questionnaire

As part of a behavioural study in which we had pioneered this paradigm^16^, a questionnaire was completed during debriefing of the study. In this study, we included all possible attentional conditions, i.e., visual-tactile, audio-visual and audio-tactile target combinations. We asked the participants to rank (1) ease of classifying the change in modalities as either increase or decrease, (2) the ease of ignoring modalities when task irrelevant and (3) ease of judging congruence for the three possible attentional conditions. Furthermore, we asked for strategies during task in open form. Statistical analysis for (1-3) was performed by means of nonparametric Wilcoxon rank sum test (ranksum.m) and correction for multiple testing was carried out according to Bonferroni. That is, reported *p*-values are multiplied with 9. Open form answers are reported anecdotally.

#### EEG: Regions of interest (ROIs) analysis

Primary cortical regions for vision, audition and tactile perception were chosen from the Freesurfer atlas which is constructed by gyral identification and parcellation based on anatomical landmarks^56^. For each frequency band, baseline-corrected, time and ROI averaged data in the change interval ([0; 300] ms relative to change onset) was evaluated by means of ANOVA with factors *ROI* (visual/auditory/somatosensory), *ATTENTION* (VT/AV) and *CONGRUENCE* (congruent/incongruent) and. Simple effects of significant ANOVA effects were assessed by paired-sample t-tests applying Bonferroni correction.

#### EEG: Whole-brain permutation statistics

Complementing ROI analysis, a whole brain exploratory analysis of differences between experimental conditions was conducted and evaluated by means of nonparametric cluster-based permutation statistics^57^. A null distribution was computed by randomly drawing trials into two sets per subject (300000 permutations). For each node of the cortical grid, a paired-sample t-test was computed between averaged, baseline-corrected power of the two sets and statistical maps were thresholded (*p* < .05). Significant clusters were found and the size of the largest cluster was noted. This procedure was carried out separately for the four frequency bands. Contrasts corresponding to a 2 (*ATTENTION*) x 2 (*CONGRUENCE*) design were computed and evaluated against the aforementioned null-hypothesis. Alpha level was adjusted according to Bonferroni (12 comparisons, α_crit_ = 0.05/12 = .0042).

Cluster statistics were complemented by post-hoc analyses that were designed (1) to detail on the time-course of the *ATTENTION* effect and (2) to disentangle the contributions of sub-conditions to the overall effect of *CONGRUENCE*.

1. For clusters showing a significant effect of *ATTENTION*, we computed the time course of average within cluster spectral power separately for visual-tactile and audio-visual conditions. Significance of the difference between time courses was evaluated using nonparametric cluster-based permutation statistics (300000 permutations). For each permutation, time courses were shuffled and paired-sample t-tests between VT and AV were computed for each sample. The number of samples included in the longest temporally continuous cluster of significant difference was noted to form the maximum statistic null distribution. In the original data, periods of significant difference between attentional conditions were considered significant in the temporal domain when they held more samples than the 99th percentile of the null distribution.
2. For this analysis we differentiated according to whether attended stimulus components were “fully congruent” or “distracted congruent”. Fully congruent (FC) means that all stimulus components, including the distracting modality, change congruently (that is, all components increased or decreased in intensity; Fig. 1b, top box). Distracted congruent (DC) means that the distractor’s change direction deviates from the change direction in the attended modalities (Fig. 1b, middle boxes). In this case, the participant has to resolve the conflict between attended congruence and unattended incongruence. In order to disentangle these two scenarios, we computed contrasts of FC respectively DC against attended incongruent conditions.

## Acknowledgements

This work was supported by grants from the German Research Foundation (SFB 936/A3 and SFB TRR 169/B1 to A.K.E. as well as DFG FR-3366-1 to U.F.), the EU (ERC-2010-AdG-269716 to A.K.E.) and the Landesforschungsförderung Hamburg (CROSS FV 25 to A.K.E.). We thank Bettina Schwab for invaluable discussions on the manuscript and Nina Noverijan for assistance in data recording.

## Author contributions

J.M., U.F. and A.K.E. designed the experiment. J.M. recorded the data. J.M. analysed the data. J.M. wrote the main manuscript text. J.M., U.F. and A.K.E. reviewed the manuscript.

## Data Availability

Behavioural and electrophysiological data will be made available upon request to the corresponding author.

## References

1. Talsma, D., Senkowski, D., Soto-Faraco, S. & Woldorff, M. G. The multifaceted interplay between attention and multisensory integration. Trends Cogn. Sci. 14, 400–410 (2010).

2. Pessoa, L., Kastner, S. & Ungerleider, L. G. Neuroimaging studies of attention: from modulation of sensory processing to top-down control. J. Neurosci. 23, 3990–3998 (2003).

3. Ling, S., Liu, T. & Carrasco, M. How spatial and feature-based attention affect the gain and tuning of population responses. Vision Res. 49, 1194–1204 (2009).

4. Peterson, E. J. & Voytek, B. Alpha oscillations control cortical gain by modulating excitatory-inhibitory background activity. bioRxiv 185074 (2017). doi:10.1101/185074

5. Worden, M. S., Foxe, J. J., Wang, N. & Simpson, G. V. Anticipatory biasing of visuospatial attention indexed by retinotopically specific alpha-band electroencephalography increases over occipital cortex. J. Neurosci. 20, RC63 (2000).

6. van Diepen, R. M., Miller, L. M., Mazaheri, A. & Geng, J. J. The role of alpha activity in spatial and feature-based attention. eNeuro 3, ENEURO.0204-16.2016 (2016).

7. Sauseng, P. et al. A shift of visual spatial attention is selectively associated with human EEG alpha activity. Eur. J. Neurosci. 22, 2917–2926 (2005).

8. Marshall, T. R., O’Shea, J., Jensen, O. & Bergmann, T. O. Frontal eye fields control attentional modulation of alpha and gamma oscillations in contralateral occipitoparietal cortex. J. Neurosci. 35, 1638–1647 (2015).

9. Siegel, M., Donner, T. H., Oostenveld, R., Fries, P. & Engel, A. K. Neuronal synchronization along the dorsal visual pathway reflects the focus of spatial attention. Neuron 60, 709–719 (2008).

10. Jensen, O. & Mazaheri, A. Shaping functional architecture by oscillatory alpha activity: Gating by inhibition. Front. Hum. Neurosci. 4, 186 (2010).

11. Bonnefond, M. & Jensen, O. Gamma activity coupled to alpha phase as a mechanism for top-down controlled gating. PLoS ONE 10, e0128667 (2015).

12. Mazaheri, A. et al. Region-specific modulations in oscillatory alpha activity serve to facilitate processing in the visual and auditory modalities. Neuroimage 87, 356–362 (2014).

13. Haegens, S., Händel, B. F. & Jensen, O. Top-down controlled alpha band activity in somatosensory areas determines behavioral performance in a discrimination task. J. Neurosci. 31, 5197–5204 (2011).

14. Schneider, T. R., Engel, A. K. & Debener, S. Multisensory identification of natural objects in a two-way crossmodal priming paradigm. Experimental Psychology 55, 121–132 (2008).

15. Göschl, F., Engel, A. K. & Friese, U. Attention modulates visual-tactile interaction in spatial pattern matching. PLoS ONE 9, e106896 (2014).

16. Misselhorn, J., Daume, J., Engel, A. K. & Friese, U. A matter of attention: Crossmodal congruence enhances and impairs performance in a novel trimodal matching paradigm. Neuropsychologia 88, 113–122 (2016).

17. Ghazanfar, A. A. & Schroeder, C. E. Is neocortex essentially multisensory? Trends Cogn. Sci. 10, 278–285 (2006).

18. Koelewijn, T., Bronkhorst, A. & Theeuwes, J. Attention and the multiple stages of multisensory integration: A review of audiovisual studies. Acta Psychologica 134, 372–384 (2010).

19. Senkowski, D., Schneider, T. R., Foxe, J. J. & Engel, A. K. Crossmodal binding through neural coherence: implications for multisensory processing. Trends Neurosci. 31, 401–409 (2008).

20. Bien, N., ten Oever, S., Goebel, R. & Sack, A. T. The sound of size: crossmodal binding in pitch-size synesthesia: a combined TMS, EEG and psychophysics study. Neuroimage 59, 663–672 (2012).

21. Schneider, T. R., Debener, S., Oostenveld, R. & Engel, A. K. Enhanced EEG gamma-band activity reflects multisensory semantic matching in visual-to-auditory object priming. NeuroImage 42, 1244–1254 (2008).

22. Konen, C. S. & Haggard, P. Multisensory parietal cortex contributes to visual enhancement of touch in humans: A single-pulse TMS study. Cereb. Cortex 24, 501–507 (2014).

23. Cappe, C. & Barone, P. Heteromodal connections supporting multisensory integration at low levels of cortical processing in the monkey. Eur. J. Neurosci. 22, 2886–2902 (2005).

24. Senkowski, D., Talsma, D., Herrmann, C. S. & Woldorff, M. G. Multisensory processing and oscillatory gamma responses: effects of spatial selective attention. Exp. Brain Res. 166, 411–426 (2005).

25. Lakatos, P. et al. The leading sense: supramodal control of neurophysiological context by attention. Neuron 64, 419–430 (2009).

26. van Atteveldt, N., Murray, M. M., Thut, G. & Schroeder, C. Multisensory integration: flexible use of general operations. Neuron 81, 1240–1253 (2014).

27. Keil, J. & Senkowski, D. Neural oscillations orchestrate multisensory processing. Neuroscientist 24, 609–626 (2018).

28. Friese, U. et al. Oscillatory brain activity during multisensory attention reflects activation, disinhibition, and cognitive control. Sci. Rep. 6, 32775 (2016).

29. Watson, A. B. & Pelli, D. G. QUEST: a Bayesian adaptive psychometric method. Percept. Psychophys. 33, 113–120 (1983).

30. Chaumon, M. & Busch, N. A. Prestimulus neural oscillations inhibit visual perception via modulation of response gain. J Cogn Neurosci 26, 2514–2529 (2014).

31. Meredith, M. A. & Stein, B. E. Visual, auditory, and somatosensory convergence on cells in superior colliculus results in multisensory integration. J. Neurophysiol. 56, 640–662 (1986).

32. Corbetta, M. et al. A common network of functional areas for attention and eye movements. Neuron 21, 761–773 (1998).

33. Grosbras, M.-H. & Paus, T. Transcranial magnetic stimulation of the human frontal eye field: effects on visual perception and attention. J. Cogn. Neurosci. 14, 1109–1120 (2002).

34. Muggleton, N. G., Juan, C.-H., Cowey, A. & Walsh, V. Human frontal eye fields and visual search. J. Neurophysiol. 89, 3340–3343 (2003).

35. Moore, T. & Armstrong, K. M. Selective gating of visual signals by microstimulation of frontal cortex. Nature 421, 370–373 (2003).

36. Silvanto, J., Lavie, N. & Walsh, V. Stimulation of the human frontal eye fields modulates sensitivity of extrastriate visual cortex. J. Neurophysiol. 96, 941–945 (2006).

37. Quentin, R., Chanes, L., Vernet, M. & Valero-Cabré, A. Fronto-parietal anatomical connections influence the modulation of conscious visual perception by high-beta frontal oscillatory activity. Cereb. Cortex 25, 2095–2101 (2015).

38. Ruff, C. C. et al. Concurrent TMS-fMRI and psychophysics reveal frontal influences on human retinotopic visual cortex. Curr. Biol. 16, 1479–1488 (2006).

39. Marshall, T. R., Bergmann, T. O. & Jensen, O. Frontoparietal structural connectivity mediates the top-down control of neuronal synchronization associated with selective attention. PLoS Biol. 13, e1002272 (2015).

40. Hecht, D. & Reiner, M. Sensory dominance in combinations of audio, visual and haptic stimuli. Exp. Brain Res. 193, 307–314 (2009).

41. Donaldson, P., Rinehart, N. J. & Enticott, P. G. Noninvasive stimulation of the temporoparietal junction: A systematic review. Neurosci. Biobehav. Rev. 55, 547–72 (2015).

42. Karnath, H. O. New insights into the functions of the superior temporal cortex. Nat. Rev. Neurosci. 2, 568–576 (2001).

43. Friedrich, F. J., Egly, R., Rafal, R. D. & Beck, D. Spatial attention deficits in humans: a comparison of superior parietal and temporal-parietal junction lesions. Neuropsychology 12, 193–207 (1998).

44. Corbetta, M. & Shulman, G. L. Control of goal-directed and stimulus-driven attention in the brain. Nat. Rev. Neurosci. 3, 201–215 (2002).

45. Göschl, F., Friese, U., Daume, J., König, P. & Engel, A. K. Oscillatory signatures of crossmodal congruence effects: An EEG investigation employing a visuotactile pattern matching paradigm. Neuroimage 116, 177–186 (2015).

46. Roa Romero, Y. et al. Alpha-band oscillations reflect altered multisensory processing of the McGurk illusion in schizophrenia. Front. Hum. Neurosci. 10, 41 (2016).

47. Wang, W., Hu, L., Cui, H., Xie, X. & Hu, Y. Spatio-temporal measures of electrophysiological correlates for behavioral multisensory enhancement during visual, auditory and somatosensory stimulation: A behavioral and ERP study. Neurosci. Bull. 29, 715–724 (2013).

48. Peng, W., Hu, L., Zhang, Z. & Hu, Y. Causality in the association between P300 and alpha event-related desynchronization. PLoS ONE 7, e34163 (2012).

49. Womelsdorf, T., Johnston, K., Vinck, M. & Everling, S. Theta-activity in anterior cingulate cortex predicts task rules and their adjustments following errors. Proc. Natl. Acad. Sci. U.S.A. 107, 5248–5253 (2010).

50. Menon, V. & Uddin, L. Q. Saliency, switching, attention and control: a network model of insula function. Brain. Struct. Funct. 214, 655–667 (2010).

51. Delorme, A. & Makeig, S. EEGLAB: an open source toolbox for analysis of single-trial EEG dynamics including independent component analysis. J. Neurosci. Methods 134, 9–21 (2004).

52. Hipp, J. F. & Siegel, M. Dissociating neuronal gamma-band activity from cranial and ocular muscle activity in EEG. Front. Hum. Neurosci. 7, 338 (2013).

53. Hassler, U., Barreto, N. T. & Gruber, T. Induced gamma band responses in human EEG after the control of miniature saccadic artifacts. Neuroimage 57, 1411–1421 (2011).

54. Pascual-Marqui, R. D. et al. Assessing interactions in the brain with exact low-resolution electromagnetic tomography. Philos. Trans. A Math. Phys. Eng. Sci. 369, 3768–3784 (2011).

55. Nolte, G. & Dassios, G. Analytic expansion of the EEG lead field for realistic volume conductors. Phys. Med. Biol. 50, 3807–3823 (2005).

56. Desikan, R. S. et al. An automated labeling system for subdividing the human cerebral cortex on MRI scans into gyral based regions of interest. NeuroImage 31, 968–980 (2006).

57. Maris, E. & Oostenveld, R. Nonparametric statistical testing of EEG- and MEG-data. J. Neurosci. Methods 164, 177–190 (2007).

